# Transcription factors across the *Escherichia coli* pangenome: a 3D perspective

**DOI:** 10.1101/2024.02.08.579464

**Authors:** Gabriel Moreno-Hagelsieb

## Abstract

**Motivation:** Identification of complete sets of transcription factors (TFs) is a foundational step in the inference of genetic regulatory networks. With the availability of high-quality predictions of protein three-dimensional structures (3D), it has become possible to use structural comparisons for the inference of homology beyond what is possible from sequence analyses alone. This work explores the potential to use predicted 3D structures for the identification of TFs in the *Escherichia coli* pangenome.

**Results:** Comparisons between predicted structures and their experimentally confirmed counterparts confirmed the high-quality of predicted structures, with most 3D structural alignments showing TM-scores well above established structural similarity thresholds, though the quality seemed slightly lower for TFs than for other proteins. As expected, structural similarity decreased with sequence similarity, though most TM-scores still remained above the structural similarity threshold. This was true regardless of the aligned structures being experimental or predicted. Results at the lowest sequence identity levels revealed potential for 3D structural comparisons to extend homology inferences below the “twilight zone” of sequence-based methods. The body of predicted 3D structures covered 99.7% of available proteins from the *E. coli* pangenome, missing only two of those matching TF domain sequence profiles. Structural analyses increased the inferred TFs in the *E. coli* pangenome by 18% above the amount obtained with sequence profiles alone.

## Introduction

Genome sequences continue to accumulate at enormous rates (Sayers *et al*., 2022; The UniProt Consortium *et al*., 2022), constantly adding to the pile of proteins to characterize. Identifying proteins within newly sequenced genomes depends mainly on the inference of homology, commonly achieved either by sequence comparison, or by comparison against databases of protein family profiles. Both methods for inferring homology are limited by the limited information from dealing with character chains.

Ultimately, protein three dimensional (3D) structures might offer the most information for the inference of homology and function (Skolnick *et al*., 2000; Leman *et al*., 2023). However, the task has been hindered by the slow accumulation of structures relative to the accumulation of sequences. The need for structures for improved clues to homology and function inspired structural genomics (Burley *et al*., 1999; Burley, 2000; Skolnick *et al*., 2000), a project aiming at solving representative structures for each inferred structure from which to predict, by homology, the structures of all the protein family members. While the project accelerated the solution of 3D structures, it still remained too slow for the pace of sequence accumulation.

Predictions of 3D structures, founded on artificial intelligence, has revolutionised the field (Varadi *et al*., 2021; Baek *et al*., 2021). The recent methods for 3D predictions have produced structures that seemingly compete with experimental 3D structure solutions (Varadi *et al*., 2021; Baek *et al*., 2021). Not only that, the Alphafold project offered to predict the structures of all proteins available at the UniProt knowledgebase (The UniProt Consortium *et al*., 2022), counting today with more than 214,000,000 3D structures (Varadi *et al*., 2023). Therefore, we count today with a very rich database for experimentation, specifically, for the theme of this work, about the inference of homology and function beyond what is possible using sequence information alone.

Because of our interest on the identification of transcription factors (TFs), proteins that play an important role in modulating gene expression, here we explore the inference of homology by 3D structural comparisons across the *Escherichia coli* pangenome. To this end, we use experimentally confirmed TFs from RegulonDB (Tierrafría *et al*., 2022), a collection of TF protein family profiles from the Pfam database (Mistry *et al*., 2020; Sanchez *et al*., 2020), as well as 3D structures both, experimental (Bittrich *et al*., 2023), and predicted (Varadi *et al*., 2023).

## Methods

### Sequence data and comparisons

Genome data were downloaded from NCBI’s RefSeq genome database (Haft *et al*., 2023) by the end of August 2023. To retrieve the *E. coli* pangenome, genome distances were estimated between all genomes classified as belonging to Enterobacterales using mash (Ondov *et al*., 2016). Genomes were clustered and cut into groups at a Mash-distance threshold of 0.048 (Hernández-Salmerón *et al*., 2023). The pangenome consisted of the group containing all the genomes labeled as *E. coli*. The identifiers of experimentally confirmed transcription factors of *E. coli* K12 MG1655, were retrieved from RegulonDB (Tierrafría *et al*., 2022).

The sequences of all experimentally available three-dimensional (3D) structures, were retrieved from the PDB (Bittrich *et al*., 2023) by the beginning of December 2023. The sequences for all Alphafold 3D structures were downloaded from the specialized database (Varadi *et al*., 2021, 2023), by February 2023 (the data has been stable since October 2022 as Alphafold version 4).

Protein sequence comparisons were run using diamond v. 2.1.8 (Buchfink *et al*., 2021), with an E-value threshold of 1e-6, and soft-masking (Hernández-Salmerón and Moreno-Hagelsieb, 2020).

### Transcription factors

To identify protein sequences of experimentally know transcription factors (TFs), we downloaded the list of TFs from *E. coli* K12 MG1655 from RegulonDB (Tierrafría *et al*., 2022). Only TFs with strong evidence were considered. Cross-checking against the genome of this strain resulted in 86 TFs sequences.

To identify further TFs, protein sequences from the *E. coli* pangenome, from the PDB, and from the SwissProt subset of Alphafold, we compared them against all available Pfam profiles (Mistry *et al*., 2020; Paysan-Lafosse *et al*., 2022) (v. 36), using MMseqs2 v. 15-6f452 (Steinegger and Söding, 2017). Pfam TFs were proteins matching Pfam domains characteristic of TF in Bacteria and Archaea (Sanchez *et al*., 2020). MMseqs2 was run with an E value threshold of 1e-3, and the results filtered for a minimum coverage of 60% of the Pfam models, with no more than 15% overlap between matched domains.

### Protein 3D structures and comparisons

Protein structures were downloaded from the PDB (Bittrich *et al*., 2023), and from Alphafold (Varadi *et al*., 2023) as necessary. Structural alignments were performed using US-align (v. 20220626) (Zhang *et al*., 2022), and foldseek (v. 8-ef4e960) (Kempen *et al*., 2023), as indicated under results. Information about clusters of Alphafold structures were downloaded from the AFDB (Barrio-Hernandez *et al*., 2023).

To divide protein 3D structures into structural domains, we used the SWORD2 pipeline with default parameters (Cretin *et al*., 2022).

## Results and Discussion

### The *E. coli* pangenome contained 2,878 genomes and 718,581 unique proteins

To define the *E. coli* pangenome, we clustered all genomes assigned to the Enterobacteriaceae taxonomic family by Mash-distances (see methods). Cutting the hierarchy at a threshold of 0.048, as suggested by previous analyses (Hernández-Salmerón *et al*., 2023), produced one group containing all but one of the genomes labeled as *E. coli*, plus all the genomes under the genus *Shigella*, as expected (Table 1) (Hernández-Salmerón *et al*., 2023). One *E. coli* genome clustered with an *Escherichia ruysiae* genome, apart from any other genome under the genus *Escherichia* or *Shigella*. Thus, we ignored this genome as a potentially mislabelled one. The resulting pangenome contained 2,878 genomes (Table 1).

**Table 1.** The *Escherichia coli* pangenome.

| Species | Count |
| --- | --- |
| *Escherichia coli* | 2703 |
| *Shigella flexneri* | 93 |
| *Shigella sonnei* | 37 |
| *Shigella dysenteriae* | 24 |
| *Shigella boydii* | 14 |
| *Escherichia* sp. | 7 |

Only seven genomes lacked a species designation.

The total number of protein sequences annotated in the *E. coli* pangenome, was 13,265,057. After filtering for identical sequences, the dataset reduced to 718,581 unique proteins.

### Pfam profiles found 31,282 unique transcription factors (TFs) in the *E. coli* pangenome

The transcription factor dataset from RegulonDB contained 86 transcription factors (TFs) with strong evidence. Cross-checking the TF names, synonyms, and annotations, against the corresponding annotations in the *E. coli* K12 MG1655 genome, allowed us to find the protein identifiers for different collections, such as SwissProt and NCBI.

To find more potential TFs in the pangenome, we compared the unique proteins dataset against all Pfam profiles using MMseqs2 (see Methods). These results were filtered to keep matches covering 60% or more of the Pfam domain profile, with no more than 15% overlap between profile matches. After filtering, 568,578, or 71.1%, proteins matched at least one Pfam domain.

Seventy-three of the 86 RegulonDB TFs matched at least one Pfam profile. Of these, 71 matched Pfam profiles typical of TFs in bacteria (Sanchez *et al*., 2020), while the other two matched one typical of Archaea (Sanchez *et al*., 2020). Therefore, to identify TFs by Pfam analysis, we included every protein matching TF Pfam domains listed as either from Bacteria or from Archaea.

The list of bacterial TF domains contained 123 Pfam identifiers, while that of archaeal TFs contained 43 (Sanchez *et al*., 2020). The union of these two lists produced 155 Pfam identifiers. A total of 31,282 unique *E. coli* proteins matched at least one of these profiles. These Pfam-inferred TFs (Pfam TFs) matched 101 of the 155 TF Pfam profiles, 19 of them found only in the list of archaeal TF pfam domains. Combined with the RegulonDB TFs, we obtained a total of 31,295 TFs.

### Alignments against experimental 3D structures confirmed the high-quality of structural predictions

To identify predicted 3D structures corresponding to experimental ones, we compared the protein sequences of the PDB against those from the Alphafold subset belonging to the model organism *E. coli* K12 MG1655 (4,363 proteins). The PDB sequences were filtered to eliminate redundant sequences from proteins in the same PDB structural file. Equivalent pairs were defined as at least 95% identical, with 95% coverage of both protein sequences in the alignment. The total number of proteins with both a PDB and an Alphafold structure was 1,319 (30.2%). Among them, 65 were Pfam TFs, and 20 were RegulonDB TFs, with 16 of the latter intersecting with the Pfam TFs.

To check the quality of Alphafold predicted 3D structures, we aligned them with their equivalent, experimental, 3D structures with US-align (see methods). Most of these alignments produced TM-scores well above the structural similarity threshold of 0.5 (Fig. 1), a threshold previously shown to correspond to structures sharing the same fold (Xu and Zhang, 2010). While also well above the structural similarity threshold, the TM-scores were lower for both, RegulonDB TFs and Pfam TFs.

**Fig. 1.**
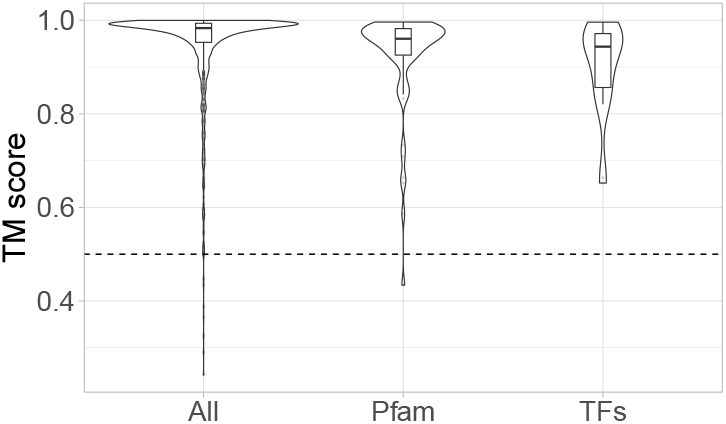
Structural alignment between *E. coli* K12 MG1655 predicted and experimental 3D protein structures. The overall quality of predicted structures was high. Though the quality for both, experimentally confirmed transcription factors (TFs), as well as proteins containing TF Pfam domains, seems lower, their structural alignments still resulted in TM scores above the 0.5 threshold, normally associated with shared structures (Xu and Zhang, 2010).

### Structural similarity between TFs decreased with sequence identity

To identify three-dimensional (3D) structures corresponding to transcription factors (TFs), we compared all PDB sequences, as well as sequences in the Alphafold subset of proteins found in SwissProt, against all Pfam profiles using MMseqs2 (see methods). As above, these results were filtered to ensure coverage of the Pfam profile equal or above 60% and no more than 15% overlap between Pfam matching segments. TFs were proteins matching Pfam domains characteristic of bacterial and archaeal TFs (Sanchez *et al*., 2020). The procedure yielded 3,901 PDB and 13,175 Alphafold/SwissProt Pfam TFs.

PDB and Alphafold Pfam TF sequences were compared against each other using diamond (see methods). Sequence alignment results were filtered to keep those showing a minimum of 95% coverage of the shortest protein in the alignment. An *ad hoc* perl script was used to select these proteins, and organise them into identity bins at 10% identity steps. The program also sampled randomly for a maximum of 250 pairs per bin. The same perl script downloaded the corresponding structures and ran pairwise structural alignments using US-align, producing a table with results that included the TM-scores obtained from the structural comparisons. The program selected the TM-scores calculated on the basis of the shortest protein.

As expected, structural similarity had a tendency to decrease with sequence divergence. Still, most results remained well above the 0.5 TM-score threshold (Fig. 2). However, results at the lowest level of sequence identity showed a tendency towards double distributions, perhaps revealing higher proportions of false positive sequence alignments. Problems at lower levels of divergence were more obvious for comparisons between predicted structures than any comparisons involving experimental ones.

**Fig. 2.**
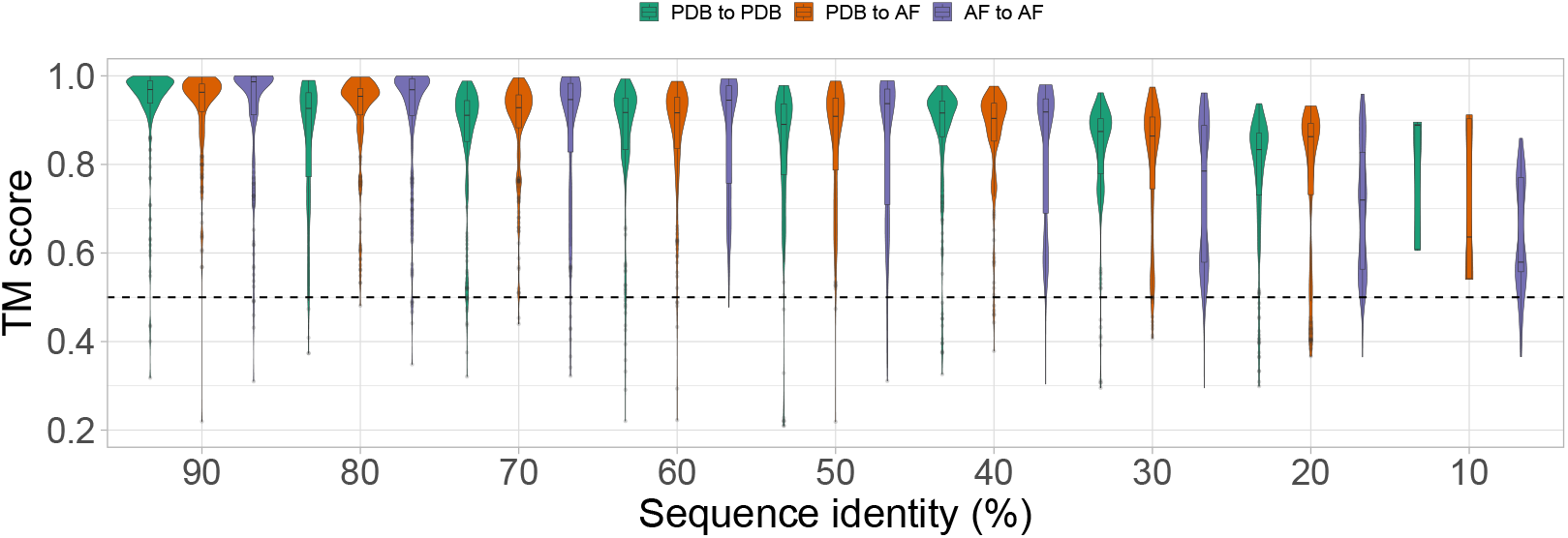
Sequence divergence *versus* structural alignments between Pfam TFs. Regardless of the compared 3D structures being experimentally determined (PDB), or predicted by Alphafold (AF), all structural alignments decreased in quality with sequence divergence, with the tendency being more obvious at the lowest sequence identity levels for comparisons between predicted structures (AF to AF). The apparent double distributions at the lowest levels of identity might indicate increased proportions of false positive sequence alignments. Still, most structural alignments had TM scores above the 0.5 threshold (dashed line), normally associated with shared structures (Xu and Zhang, 2010).

### Alphafold matched 99.7% proteins of the *E. coli* pangenome

After sequence comparisons, 718,581 unique proteins of the *E. coli* pangenome, 373,702 (52%) found matches against proteins with experimentally determined 3D structures (PDB). Of these, only 150,343, 40.2% (or 20.1% of the total) were “90/90” alignments, consisting of those with identities above 90%, as well as sequence coverage of the *E. coli* protein above 90% (Fig. 3, Table 2). Overall PDB matches across the *E. coli* pangenome ranged from 55.1% to 68.2%, with a median of 62.2%.

**Table 2.** Transcription factor inference.

| Dataset | Aligned | 90/90 | Pfam(90/90) | Structural(90/90) |
| --- | --- | --- | --- | --- |
| Pangenome | 716,730 | 693,862 | 30,997 | 36,645 |
| Alphafold | 348,386 | 340,753 | 14,731 | 17,458 |

**Fig. 3.**
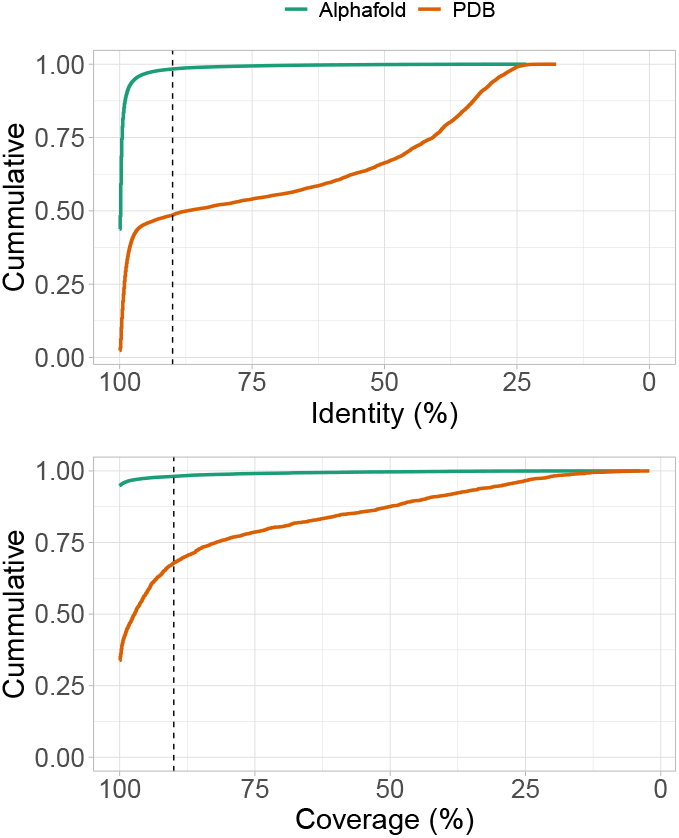
Statistics for best sequence alignment matches. Predicted (Alphafold) structures had much better sequence-based matches against the unique protein dataset annotated in the *E. coli* pangenome, than experimental (PDB) ones. More than 95% of Alphafold matches had sequence coverage and identity values above 90% (dashed lines)

When compared against the protein sequences of predicted 3D structures (Alphafold), 716,730, or 99.7%, of the unique proteins found a match. Of these, 693,862, or 96.8% (96.6% of the total), constituted 90/90 matches (Fig. 3, Table 2). Overall, Alphafold structures matched between 91.8% and 99.4% of the proteins annotated in the genomes analysed, with a median of 99.1%. It thus seems unlikely that Alphafold missed proteins of wide interest for the *E. coli* scientific community.

PDB matched 19,557 of the 31,295 protein in the unique TF dataset (62.5%), while Alphafold matched all but two of the unique TFs (proteins matching a Pfam TF domain). The two missing TFs were found each in a single one of the 2,878 genomes in the dataset. The 90/90 Alphafold proteins matched 31,010 of the unique TFs (99.1%). Thus, Alphafold should provide enough material to try and infer further TFs by structural similarity.

### *E. coli* TFs clustered into 710 structural families

To find structural clusters, or structural families, where the unique *E. coli* TFs belonged, we found the 90/90 identifiers of the Alphafold sequences matching unique TFs. The 31,010 unique TFs with 90/90 results matched 14,744 Alphafold sequences (more than one TF matched the same Alphafold sequence). AFDB is a database containing clusters of Alphafold structures (Barrio-Hernandez *et al*., 2023), based on alignments performed with the foldseek fast 3D structural alignment program (Kempen *et al*., 2023). We therefore cross-checked the 14,744 90/90 Alphafold TF identifiers against the cluster information found at the AFDB. These 14,744 belonged to 719 Alphafold clusters, nine of them with no other members in the group (singletons). Thus we found 710 structural families.

### AFDB TFs identified 18% additional TFs compared to Pfam

The complete set of structure identifiers within the 710 structural families identified above, represented a total of 5,183,897 protein structures. With this list, we checked back the unique *E. coli* proteins *versus* Alphafold sequence alignment results. This way we identified 17,615 Alphafold structures, corresponding to the 710 TF structural families found above. Most of these, 17,458 (99.1%), matched 90/90 alignment results (Table 2). Among these 17,458 AFDB TFs, 14,731 matched a Pfam profile, leaving 2,727 potential TFs discovered by structural alignment (18.5% above the 14,731 that would be found by Pfam analysis alone). Overall, these 17,458 AFDB TFs matched 36,645 of the unique *E. coli* proteins. Of these, 30,997 matched a Pfam TF domain. Thus, if these AFDB clusters are actually composed of TFs, structural comparisons increased TF prediction by 18.2% compared to Pfam profiles in the *E. coli* pangenome.

Further inspection of the AFDB TFs showed that three of the eleven RegulonDB TFs that did not match a Pfam TF profile, belonged to structural groups with proteins that previously matched a Pfam TF profile, suggesting that structural alignments can add members to previously described protein families.

UniProt runs a battery of analyses to try and assign functions to most of the protein sequences in the “knowledgebase” (The UniProt Consortium *et al*., 2022). Since the Alphafold 3D structure collection consists of predicted structures for a full dataset of UniProt proteins (Varadi *et al*., 2023), finding the corresponding annotations is straightforward, and might help us estimate the quality of TFs added by the AFDB analyses above.

In the case of Alphafold proteins matching Pfam TFs, we found UniProt annotations for 7,991 of the 14,731 proteins. Among these, 5,729, or 71.7%) matched Gene Ontology (GO) (Aleksander *et al*., 2023) keys found in the RegulonDB TFs, whose descriptions match TF activities (GO keys proper of TF activities).

Of the extra 2,727 Alphafold proteins found by AFDB analyses, potentially new TFs that did not match a Pfam TF profile, 1,474 had UniProt annotations. Among these, 456, or 30.9%, matched GO keys proper of TF activities. This result suggests that there is room for 3D structural comparisons for improving over UniProt’s battery of analyses.

Structural families might structurally align over domains other than the TF domains. Therefore, to ensure that the 2,842 potential TF structures found by our AFDB analyses had a TF domain, we produced structural alignments between the ones matching Pfam TF profiles against the 2,727 that did not match a Pfam TF profile using foldseek (Kempen *et al*., 2023). We then checked if the coordinates of the query protein matching the Pfam TF profiles were covered, in the structural alignment, by the target proteins (Fig. 4). This way we confirmed that 1,914 (70.2%) of the 2,727 Alphafold proteins that did not match a Pfam TF profile, aligned, structurally, with regions corresponding to such profiles in the query proteins. These matches considered only structural alignments with TM-scores above 0.5, that covered completely the regions matching a Pfam TF profile. The remaining proteins also covered these regions, but the TM-scores were below 0.5.

**Fig. 4.**
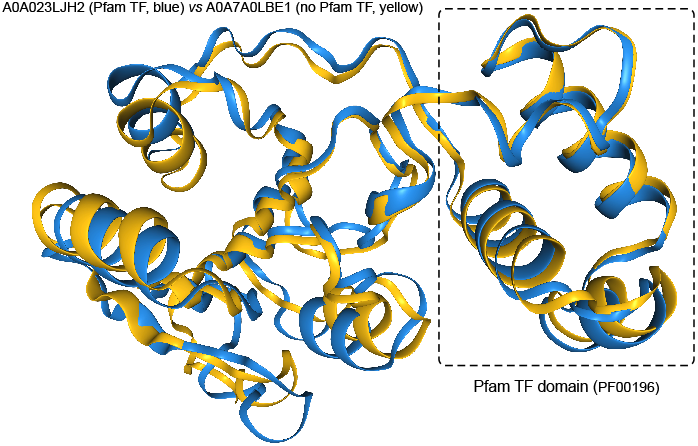
Structural inference of transcription factors. Comparing structures whose sequences matched Pfam domains, against structures lacking such matches, while ensuring that the structural alignments covered the appropriate segments, allowed for the inference of transcription factors by structural similarity

## Conclusion

In this work we have confirmed the high-quality of 3D structures predicted by Alphafold. We have also found that, though 3D structural alignments are less reliable for predicted structures than for experimental ones, when the proteins have low sequence identity, they still offer potential for the inference of homology beyond what is possible using sequence family profiles. Our results allowed us to increase our database of inferred TFs in the *E. coli* pangenome by 18%. Further research might be necessary to confirm the quality of these inferences.

## Author contributions statement

The whole enchilada: GMH

## Acknowledgments

This work was supported by a Discovery Grant (RGPIN-2018-06180) awarded to GMH by the Natural Sciences and Engineering Research Council (NSERC) of Canada.

